# Transcriptome responses of the aphid vector *Myzus persicae* are shaped by identities of the host plant and the virus

**DOI:** 10.1101/2022.07.18.500449

**Authors:** Quentin Chesnais, Victor Golyaev, Amandine Velt, Camille Rustenholz, Maxime Verdier, Véronique Brault, Mikhail M. Pooggin, Martin Drucker

## Abstract

**Background:** Numerous studies have documented modifications in vector orientation behavior, settling and feeding behavior, and/or fecundity and survival due to virus infection in host plants. These alterations are often expected to enhance virus transmission, which has led to the hypothesis that such effects are vector manipulations by the virus. However, until now, the gene expression changes correlating with these effects and indicative of modified vector pathways and mechanisms are mostly unknown.

**Results:** Transcriptome profiling of *Myzus persicae* aphids feeding on turnip yellows virus (TuYV) and cauliflower mosaic virus (CaMV) infected *Arabidopsis thaliana* and *Camelina sativa* revealed a substantial proportion of commonly deregulated genes, amongst them many with general functions in plant-virus-aphid interactions. We identified also aphid genes specifically deregulated by CaMV or TuYV infection, which might be related to the viral transmission mode. Furthermore, we observed strong host-specific differences in the gene expression patterns with plant virus infection causing more deregulations of aphid genes on *A. thaliana* than on *C. sativa*, likely related to the differences in susceptibility of the plant hosts to these viruses. Finally, stress-related aphid genes were downregulated in *M. persicae* on both infected plants, regardless of the virus.

**Conclusions:** TuYV, relying on the circulative persistent mode of transmission, tended to affect developmental genes. This could increase the proportion of alate aphids, but also affect their locomotion, neuronal activity, and lifespan. CaMV, using the non-circulative non-persistent mode of transmission, had a strong impact on feeding-related genes and in particular those related to salivary proteins. In general, these transcriptome alterations targeted pathways that seem to be particularly adapted to the transmission mode of the corresponding virus and could be evidence of vector manipulation by the virus.

## Introduction

Aphids are major pests not only because they deprive plants of nutrient resources when feeding on phloem sap but also because they transmit many plant-pathogenic viruses. Indeed, most plant viruses rely on vectors for transmission to a new susceptible host (Dietzgen et al., 2016). As phloem-feeders, aphids play a preponderant role in plant virus transmission, because their particular feeding behavior allows direct delivery of virus particles into the cytoplasm of cells in the epidermis, mesophyll, vascular tissue and/or the phloem sap of a new host. Their hypodermic needle-like mouthparts, the stylets, can penetrate cuticle and cell walls and enter into plant cells and sieve tubes without inflicting any major damage. More precisely, aphids alighting on a new plant will initiate probing phases (i.e. test the potential host for suitability) consisting of extracellular pathways and exploratory intracellular punctures into the epidermis and underlying tissues and, if accepted, plunge then their stylets into the sieve cells whose sap constitutes their principal food source (Tjallingii & Hogen Esch, 1993). During the probing and feeding phases, aphids secrete different saliva types that contain amongst other compounds effector molecules modulating interactions with the plant immune system and susceptibility (Rodriguez & Bos, 2013).

Viral infection often modifies plant phenotypical traits such as leaf color, morphology, surface properties, composition and quantity of volatile organic compounds (VOCs) and metabolites (Matthews, 2014). This may impact vector behavior and performance, i.e. attract/deter vectors, modify their feeding behavior and incite/discourage colonization (reviewed by Fereres & Moreno, 2009). There is evidence that such virus-mediated modifications can facilitate virus transmission, a concept known as ‘pathogen manipulation’. The modifications depend on the viruses’ modes of transmission (Mauck et al., 2012, 2018). So-called non-persistent/non-circulative viruses have fast transmission kinetics and are acquired and inoculated, but also lost from the vectors, within seconds to minutes (Day & Irzykiewicz, 1954). The non-circulative viruses rely on other parameters for optimal transmission than so-called persistent/circulative viruses that have slow transmission kinetics (hours to days) as the virus is injected as a saliva component into a new host, following the passage of viral particles from the intestine to the salivary glands through the hemolymph. Consequently, vectors retain and transmit the circulative viruses for weeks or lifelong (reviewed by Gray & Banerjee, 1999).

How virus-mediated plant modifications translate into changes in vector behavior and performance, is largely unknown (reviewed by Fereres & Moreno, 2009; Dáder et al., 2017; Mauck et al., 2019). It is assumed that most of the modifications are indirect, i.e. aphids and other vectors react to virus-induced changes in the plant. For example, yellowing symptoms induced by virus infection may attract and encourage the settling of insect vectors (for example, Johnston & Martini, 2020; Chesnais et al., 2022). A well-characterized example of such plant modifications by non-persistent, non-circulative viruses is the cucumber mosaic virus (CMV, genus *Cucumovirus*, family *Bromoviridae*). VOCs emitted by CMV-infected squash attract the green peach aphid (*Myzus persicae*, hereafter Myzus), but once landed on the infected squash, the poor palatability of the plant incites the aphids to leave fast (Mauck et al., 2010, 2014). This aphid behavior is perfectly adapted to an efficient acquisition and transmission of CMV which relies on a short acquisition time and a rapid dispersal for propagation (Bhargava, 1951). Therefore, this example might be considered as ‘host manipulation’. Although there are more examples in the literature (reviewed by Fereres & Moreno, 2009; Dáder et al., 2017), host-induced vector manipulation by non-persistent/non-circulative viruses is rather under-explored. The non-circulative virus studied here – cauliflower mosaic virus (CaMV, genus *Caulimovirus*, family *Caulimoviridae*) – follows the same transmission kinetics as CMV except that it is retained longer (hours range) in its aphid vectors (Markham et al., 1987) and therefore its transmission mode has been classified also as ‘semi-persistent’, a term coined by Sylvester (1956). Previous work (Chesnais et al., 2019) showed that Myzus vectors did not show any preference for *Camelina sativa* (hereafter Camelina) plants infected with the severe CaMV isolate B-JI, but the number of intracellular probing punctures was increased and phloem ingestion and fecundity reduced on infected plants. Using CaMV-infected *Arabidopsis thaliana* (hereafter Arabidopsis) as virus host, Myzus spent less time in the pathway phase and more time feeding on phloem and aphid fecundity was lowered, compared to healthy control plants (Chesnais et al., 2021). A similar feeding behavior was observed for Myzus feeding on Arabidopsis infected with the milder CaMV isolate Cm1841r but in contrast to plants infected with the B-JI isolate, fecundity was not affected (Chesnais et al., 2021). Thus, there are contrasting results on possible manipulation of host plants by CaMV that might depend on the virus isolate and host plant species.

Quite a body of evidence for ‘manipulation’ by persistent/circulative viruses has been collected for poleroviruses (genus *Polerovirus*, family *Solemoviridae*). Most studies on poleroviruses have shown that virus-infected plants are more attractive to aphids than healthy plants and that aphid feeding is improved and fecundity higher on infected plants (reviewed by Bosque-Pérez & Eigenbrode, 2011; Dáder et al., 2017; Mauck et al., 2018). Curiously, aphid preference changed after polerovirus acquisition, and aphids carrying poleroviruses preferred healthy plants over virus-infected plants (Alvarez et al., 2007; Carmo-Sousa et al., 2016). There is evidence that purified virus particles can bring along this preference change (Ingwell et al., 2012), indicating that for persistent/circulative viruses, not only host plant-mediated changes but also direct virus-mediated changes in aphids are to be considered. The circulative virus studied here, turnip yellows virus (TuYV, genus *Polerovirus*, family *Solemoviridae*), increases emission of VOCs in two host plants, Arabidopsis and Camelina, but only TuYV-infected Camelina, and not TuYV-infected Arabidopsis, attracted Myzus more than did healthy control plants (Claudel et al., 2018). Aphids feed longer from the phloem of TuYV-infected Camelina than from that of healthy Camelina, which might favor the acquisition of phloem-limited TuYV (Chesnais et al., 2019). Recently, a post-acquisition effect of TuYV was observed: virus-carrying Myzus aphids showed increased vector locomotory and fecundity as well as prolonged phloem feeding behavior. However, in this study, the authors did not distinguish between direct effects of the virus on the vector and indirect effects mediated by the infected host plant (Chesnais et al., 2020).

While virus-mediated effects on aphids and other hemipteran vectors are well documented, knowledge on the molecular mechanisms and the involved aphid genes is scarce. Published examples indicate that deregulation of aphid genes related to stress, cuticle, development and nucleic factors is a common feature of aphids feeding on plants infected with poleroviruses or luteoviruses (Brault et al., 2010; Li et al., 2020; Patton et al., 2021). For non-circulative viruses, the effect of viral infection of plants on aphids seems more variable. CMV acquisition by Myzus from infected tobacco changed the expression of vector genes related to metabolism, stress, and cuticle (Liang et al., 2021), whereas a study on the soybean aphid *Aphis glycines* fed on soybean plants infected with soybean mosaic virus (SMW, genus *Potyvirus*, family *Potyviridae*) has revealed only minor changes in aphid gene expression (Cassone et al., 2014).

In this paper, we explored how infection of plants with circulative versus non-circulative virus affects the transcriptome of viruliferous aphids. We specifically addressed whether the transmission mode influenced the aphid transcriptome profiles and whether alterations in the aphid transcriptome correlated with distinct behaviors of viruliferous aphids. We identified common and virus-specific deregulated genes as well as plant host-specific effects on aphids. The aphid *M. persicae* was selected for this study because it is an excellent vector for both the circulative, persistent TuYV and the non-circulative, semi-persistent CaMV. On the plant side, we selected two species of the same family *Brassicaceae*, *A. thaliana* and *C. sativa* that are suitable hosts for both viruses.

## Material and methods

### Aphids

The green peach aphid (*Myzus persicae* Sulzer, 1776) clone WMp2, originally isolated in the Netherlands (Reinink et al., 1989) and maintained in Colmar since 1992, was used for the experiments. It was reared on Chinese cabbage (*Brassica rapa* L. *pekinensis* var. Granaat) in a growth chamber at 20±1 °C and a 16 h photoperiod. Plants were grown in TS 3 fine substrate (Klasmann-Deilmann) in round 13 cm diameter pots and watered with fertilizer 209 (Fertil SAS) dissolved in tap water. Only wingless morphs were used in assays. For synchronization, adults were placed on detached Chinese cabbage leaves that were laid on 1 % agarose (Euromedex) in a Petri dish. The adults were removed 24 h later and the newborn larvae used in transcriptomic experiments 5 days later.

### Viruses

CaMV isolate Cm1841r (Chesnais et al., 2021), which is an aphid-transmissible derivative of isolate Cm1841 (Tsuge et al., 1994), and TuYV isolate TuYV-FL1 (Veidt et al., 1988) were maintained in Arabidopsis Col-0 and propagated by aphid inoculation of 2-week-old plants. Plant growth conditions were as described below.

### Virus infection and aphid infestation

Seeds of *Arabidopsis thaliana* Col-0 or *Camelina sativa* var. Celine were germinated in TS 3 fine substrate (Klasmann-Deilmann) in 7*7 cm pots and watered with tap water. Growth conditions were 14 h day 10 h night with LED illumination and a constant temperature of 21±1 °C. Two-week-old plants were inoculated with 3-5 wingless Myzus aphids that had been allowed a 24 h acquisition access period on Arabidopsis infected with TuYV or CaMV or on healthy Arabidopsis. Plants were individually wrapped in clear plastic vented bread bags to prevent cross contamination. Aphids were manually removed after a 48 h inoculation period. Eighteen days post-inoculation (dpi), 25 to 30 synchronized 5-day-old non-viruliferous aphids were placed for infestation on the rosette (Arabidopsis) or the apical leaves (Camelina) of CaMV- or TuYV-infected or mock-inoculated plants. After 72 h infestation (= 21 dpi), aphids were collected with a brush. Three biological replicates were used for analysis. For Arabidopsis, one biological replicate consisted of 4 plants, from which 25-30 aphids were collected (total of 100-120 aphids). For Camelina, one replicate was 3 plants from which 30 aphids were collected (total of 90-100 aphids). Aphid samples were deep-frozen by placing them in a −80 °C freezer and conserved at this temperature until processing.

### RNA purification and Illumina sequencing

Total RNA was extracted from aphids with TRI Reagent (Molecular Research Center) and chloroform followed by isopropanol (Merck) precipitation. Briefly, 10-50 mg of frozen aphids were placed in a mortar cooled with liquid N2, homogenized in 1 ml TRI Reagent and incubated for 2 hours at room temperature. Subsequent phase separation by addition of 200 μl cold chloroform (Merck) and centrifugation for 15 min at 12,000 g was followed by RNA precipitation with 500 μl cold isopropanol. After 10 min centrifugation at 12,000 g, the RNA pellet was washed twice with 1 ml of 75 % ethanol (Merck), air-dried and resuspended in 30 μl RNase-free water (Merck). RNA quantity and purity were measured using NanoDrop 2000c Spectrophotometer (Thermo Fisher Scientific). RNA integrity was verified by capillary electrophoresis on LabChip GX (Perkin Elmer).

Illumina sequencing of 18 aphid total RNA samples was performed at Fasteris (www.fasteris.com) using a standard protocol with the TruSeq Stranded mRNA Library Prep kit (Illumina). All the libraries (3 biological replicates per each of the six conditions [i.e., aphids on mock-inoculated, TuYV- and CaMV- infected Arabidopsis and aphids on mock-inoculated, TuYV- and CaMV- infected Camelina]) were multiplexed in one NovaSeq flowcell SP-200 with 2×75 nt paired-end customized run mode. The resulting 75 nt reads from each library were used for Myzus transcriptome profiling.

### RT-qPCR

For RT-qPCR analysis of Myzus gene expression, 10 μg total RNA was converted into cDNA using AMV Reverse Transcriptase (Promega) and oligo-dT primer. Real-time qPCR reactions (10 μl) including 3 μl of cDNA, 0.5 μl of each 10 μM primer, 5 μl of SybrGreen master mix (Roche) and 1 μl of water were processed in a LightCycler^®^ 480 instrument (Roche) using the SybrGreen master mix (Roche) following the recommended protocol. The thermocycler conditions were as follows: pre-incubation at 95 °C for 5 min, followed by 40 cycles of 95 °C for 10 s, 58 °C for 20 s and 72 °C for 20 s. The gene expression was normalized to the Myzus persicae internal reference gene EF1alpha (Naessens et al., 2015; Webster et al., 2018) (for primer sequences of targeted genes and of the internal reference gene see Table S2).

### Raw data processing and quality control for transcriptome profiling

Processing was carried out on the Galaxy France platform (https://usegalaxy.fr/) (Afgan et al., 2016). Raw reads quality was checked with FastQC (v0.11.8) and the results were then aggregated with MultiQC (v1.9). Reads were aligned on the *Myzus persicae* O reference genome (‘Myzus_persicae_O_v2.0.scaffolds.fa’ (annotations: ‘Myzus_persicae_O_v2.0.scaffolds.braker2.gff3’) downloaded from BIPAA portal (https://bipaa.genouest.org/sp/myzus_persicae/) (Mathers et al., 2021) with STAR (v2.7.6a) and quality was again checked with MultiQC. Gene counts were obtained with featureCounts (v2.0.1). Differential gene expression was then analyzed with SARTools (v1.7.3) and the DESeq2 method (i.e. aphids on TuYV-infected plants vs. mock-inoculated plants, aphids on CaMV-infected plants vs. mock-inoculated plants, and aphids on TuYV-infected plants vs. CaMV-infected plants). GO enrichment analysis of the DEGs was performed with GOseq (v1.36.0) (Young et al., 2010). We used default parameters for all steps except for featureCounts where the following deviating parameters were used: only the primary alignment was taken into account (not multi-mapped reads), exclude chimeric fragments = Yes (-C option, signifying that the fragments that have their two ends aligned to different chromosomes were NOT included for summarization), minimum base overlap = 1.

## Results and discussion

### Quality control and validation of RNA-seq data

For aphids on Arabidopsis, between 64.6 and 88.8 million 75 nt paired-end reads were obtained with a mean phred score >30 for all bases. For aphids on Camelina, between 61.8 and 82.4 million 75 nt paired-end reads were obtained with a mean phred score >30 for all bases. In all samples, there were no overrepresented sequences and only a few adapter-containing reads (0.20 % reads with adapter sequence at the last bases). Between 85.6 % and 88.7 % of reads were uniquely mapped to the aphid genome for Arabidopsis (Supplementary Table S1a) and between 81.8 % and 87.6 % of reads were uniquely mapped to the aphid genome for Camelina (Supplementary Table S1b). Of these, between 87.4 % and 88.3 % of uniquely aligned reads were assigned to an aphid gene on Arabidopsis and 83.4 % to 86.8 % aligned reads were assigned to an aphid gene on Camelina. We did not look for the nature of the unaligned reads; they might derive from endosymbionts, contaminating biologic material from plants, fungi, bacteria and the like.

Exemplarily, a similar trend of gene deregulation was confirmed by RT-qPCR for four Myzus genes with different levels of deregulation and expression, but the same trend in both infection conditions. We screened only four genes as in a previous work on aphid transcriptomics (Liu et al., 2012). Three genes showed the same trend of downregulation in RNA-seq and RT-qPCR experiments, while the forth (g15329) was found to be upregulated in all RNA-seq and RT-qPCR experiments, except for RT-qPCR on TuYV-infected plants (Supplementary Figure S1). The discrepancy in the results for g15329 expression was likely due its weak expression changes that in general are difficult to detect by RT-qPCR because of the exponential amplification kinetics of this technique. We observed the same phenomenon in a previous validation experiment (Chesnais, Golyaev, et al., 2022).

Principal component analysis of RNA-seq datasets (Figure 1a) indicated good clustering of the three biological replicates of aphids fed on mock-inoculated or virus-infected Arabidopsis. One of the three biological replicates of aphids fed on CaMV-infected Arabidopsis grouped less well with the other two but was still within an acceptable range. In the case of Camelina, the three replicates for each virus (TuYV and CaMV) clustered well together, indicating homogeneity of the replicates (Figure 1b). The Myzus data for Camelina infected with TuYV and CaMV were more similar to each other than those for Arabidopsis, indicating that the transcriptome changes in aphids fed on Camelina were less dependent on the virus species than those on Arabidopsis. In the case of mock-inoculated Camelina, two of the three replicates clustered together and were well separated from the data for virus-infected Camelina, while the third replicate clustered with the aphid data from infected plants and was therefore excluded from further analysis (Figure 1b). The transcriptomes of the plants used here for aphid infestation were analyzed in another study (Chesnais, Golyaev, et al., 2022). There the three replicates from mock-infected Camelina clustered closely together in principal component analysis. This indicates that the outlier behavior observed here was not caused by the plant itself but by another cause, which remains elusive. Taken together, all samples except one mock replicate of Camelina were suitable for transcriptome analysis.

**Figure 1.**
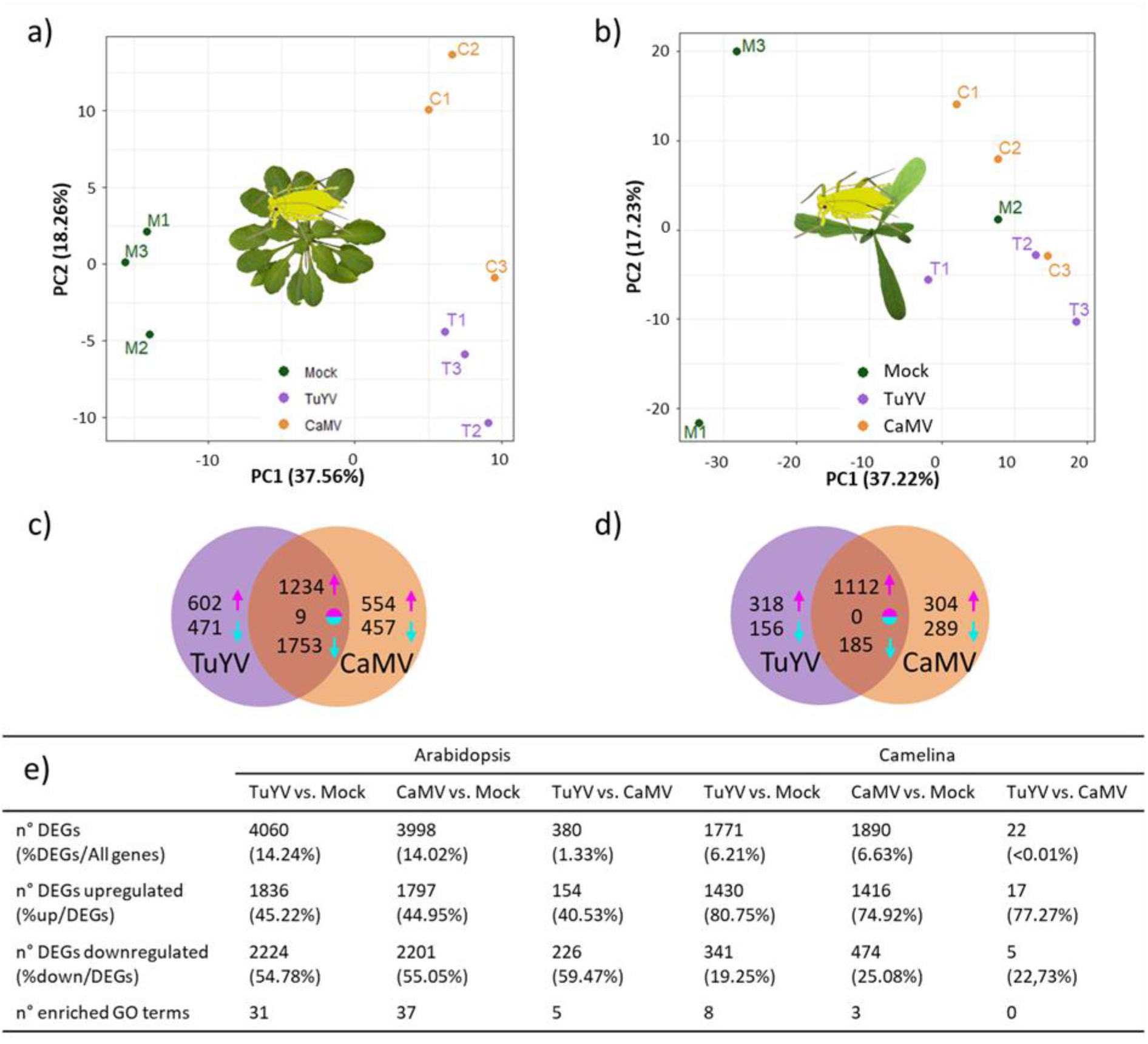
Analysis of the transcriptome profiles of aphids fed on mock-inoculated vs TuYV- and CaMV-infected plants. (a-b) Principal component analysis of three biological replicates for each condition of *Myzus persicae* feeding on (a) *Arabidopsis thaliana* and (b) *Camelina sativa*. The dots of the same color correspond to the biological replicates for each condition. The mock 2 (M2) Camelina sample was excluded from the analysis because it did not cluster with the other two replicates. (c-d) Venn diagrams presenting the number of differentially expressed genes (DEGs) in aphids fed on TuYV- and CaMV-infected Arabidopsis (c) and Camelina (d), compared to respective mock-inoculated controls. Magenta arrows: number of up-regulated genes, cyan arrows: number of down-regulated genes and two-color circles: inversely regulated genes (up-regulated genes in one virus-infected modality and down-regulated in the other virus-infected modality). e) The number of DEGs and enriched GO categories in aphids fed on TuYV and CaMV-infected plants vs mock controls as well as on TuYV- vs CaMV-infected plants.

### Global analysis of differentially expressed aphid genes

Analysis of RNA-seq data revealed twice as many differentially expressed genes (DEGs) (false discovery rate <0.05) in aphids feeding on virus-infected Arabidopsis (4,060 for TuYV and 3,998 for CaMV) as in aphids feeding on virus-infected Camelina (1,771 for TuYV and 1,890 for CaMV), compared to aphids from mock-inoculated controls (Figure 1c, 1d and 1e). Remarkably, each virus modified the expression of about the same number of genes in aphids fed on the same host. Moreover, for each plant species, about 2/3 of aphid DEGs were common for the two viruses, indicating a profound common response of aphids to feeding on infected plants, independent of the virus species and of the transmission mode (Figure 1c and 1d). Like the number of aphids DEGs, also the proportion of up- and downregulated aphid genes was virus-independent, with ca. 45 % of the aphid DEGs being upregulated after feeding on TuYV- or CaMV-infected Arabidopsis, and ca. 75 % and ca. 81 % being upregulated after feeding on TuYV- and CaMV-infected Camelina, respectively (Figure 1e). The differences in the number and proportion of up- and downregulated aphid DEGs between Arabidopsis and Camelina indicated an important plant species effect on the aphid transcriptome, which was independent of the virus. On the other hand, for each plant species, ca. 1/3 of the aphid DEGs was specific for each virus, indicating that the virus species (and, possibly, the transmission mode) had a substantial and characteristic impact on the aphid transcriptome.

### Impact of CaMV and TuYV infection on aphid metabolic pathways

#### Gene ontology analysis of Myzus infesting CaMV- or TuYV-infected plants

Using gene ontology (GO) analysis, we first looked at the effects of virus-infected Arabidopsis on the aphid transcriptome (Figure 2). In aphids fed on TuYV-infected Arabidopsis, 11 of the Top 25 enriched GO categories of DEGs classified as Biological Processes (BP) (Figure 2a). The most affected processes were ‘oxidation-reduction’ (BP), ‘integral component of membrane’ (belonging to the category Cellular Component [CC]), and the rather general process ‘ATP-binding’ (belonging to the category Molecular Function [MF]). Other prominent processes were related to protein synthesis and metabolism (translation initiation, protein synthesis, endopeptidase activity, protein folding, proteasome-mediated protein degradation and unfolded protein binding). Similarly, the most deregulated processes of aphids feeding on CaMV-infected Arabidopsis were ‘oxidation-reduction (BP)’, ‘integral component of membrane (CC)’ and ‘ATP binding (MF)’, followed by protein synthesis and metabolism-related processes (Figure 2b).

**Figure 2.**
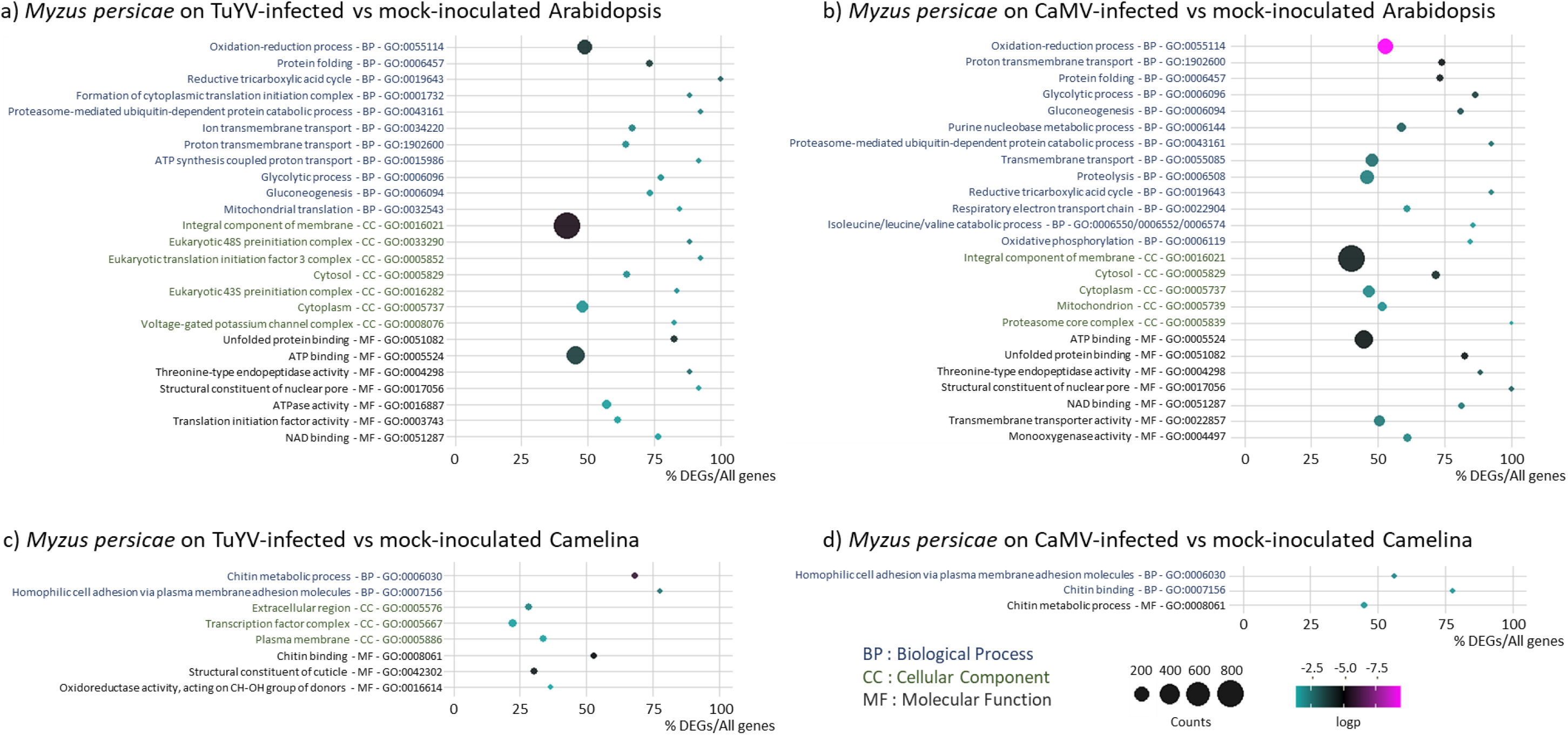
Gene ontology (GO) analysis of differentially expressed genes in *Myzus persicae* feeding on TuYV- and CaMV-infected Arabidopsis and Camelina. a) *Myzus persicae* on TuYV-infected vs mock-inoculated Arabidopsis, b) *Myzus persicae* on CaMV-infected vs mock-inoculated Arabidopsis, c) *Myzus persicae* on TuYV-infected vs mock-inoculated Camelina, and d) *Myzus persicae* on CaMV-infected vs mock-inoculated Camelina. The deregulated processes and the corresponding GO categories and IDs are specified in the vertical axis. For each GO category (BP: Biological Process, CC: Cellular Component, and MF: Molecular Function), the GO terms/processes are sorted according to decreasing log2 (1/p-value), also indicated by the color of each spot, to place the most significantly enriched GOs on top of the graph. The absolute number of DEGs that matched the GO term is indicated by the size of each spot, whereas the horizontal axis shows the percentage of DEGs belonging to the GO term.

A different picture was found for Myzus on virus-infected Camelina (Figure 2c). In the case of TuYV infection, only 8 categories [2 in biological processes (BP), 3 in cellular components (CC) and 3 in molecular functions (MF)] were identified by GO Top 25 analysis as being significantly enriched. Three of them (Figure 2d) were also identified in aphids from CaMV-infected Camelina, but none of them in aphids from infected Arabidopsis. The enriched processes included chitin-related processes (chitin binding, MF; chitin metabolic processes, BP; structural constituent of cuticle, MF), transcription (transcription factor complex, CC), oxidation reduction (oxidoreductase activity, MF) and plasma membrane-related processes (homophilic cell adhesion via plasma membrane, BP; plasma membrane, CC; extracellular region, CC). Although none of these GOs figured among the Arabidopsis Top 25 GO, there were three GO categories (related to oxidation/reduction and plasma membrane processes) that were similar to GOs identified in aphids fed on Arabidopsis.

Taken together, GO analysis revealed distinct, plant host-specific impacts on the aphid gene expression, which were rather independent of the virus species. Feeding on virus-infected Arabidopsis had a much more profound impact on aphids than feeding on virus-infected Camelina (Figure 2a,b vs 2c,d).

Since the current annotation of the Myzus genome was not as advanced as for other model organisms such as *Drosophila melanogaster*, we complemented our above-described GO analysis by analysis of Kyoto Encyclopedia of Genes and Genomes (KEGG) pathways (Kanehisa, 1996). This analysis showed that similar percentages of genes involved in ‘genetic information processing’, ‘metabolism’ and ‘signaling and cellular processes’ were modified in aphids feeding on both plant hosts (Figure S3).

### General heatmap analysis of DEGs

To better visualize the aphid transcriptome changes, heatmaps presenting all DEGs in aphids infesting Arabidopsis or Camelina were generated (Figure 3a and 3b). The profiles of all aphid replicates fed on mock-inoculated Arabidopsis or Camelina clustered well together, while the profiles from aphids feeding on virus-infected plants (CaMV and TuYV) did not. This again indicated that, in our experiment, effects of infection on aphid genes were largely independent of the virus species. Like the global analysis (Figure 1e), the heat maps showed also that Myzus on virus (TuYV or CaMV)-infected Camelina displayed proportionally more up- than downregulated genes, compared to Myzus on mock-inoculated Camelina, whereas proportions of up- and downregulated DEGs in aphids feeding on virus-infected vs mock-inoculated Arabidopsis were similar. The significance of these plant host-specific effects remains to be investigated. We speculate that Myzus might have more difficulties in establishing infestation on Arabidopsis than on Camelina, visible by the higher number of DEGs that is indicative of extensive transcriptome reprograming to adapt to the new plant host.

**Figure 3.**
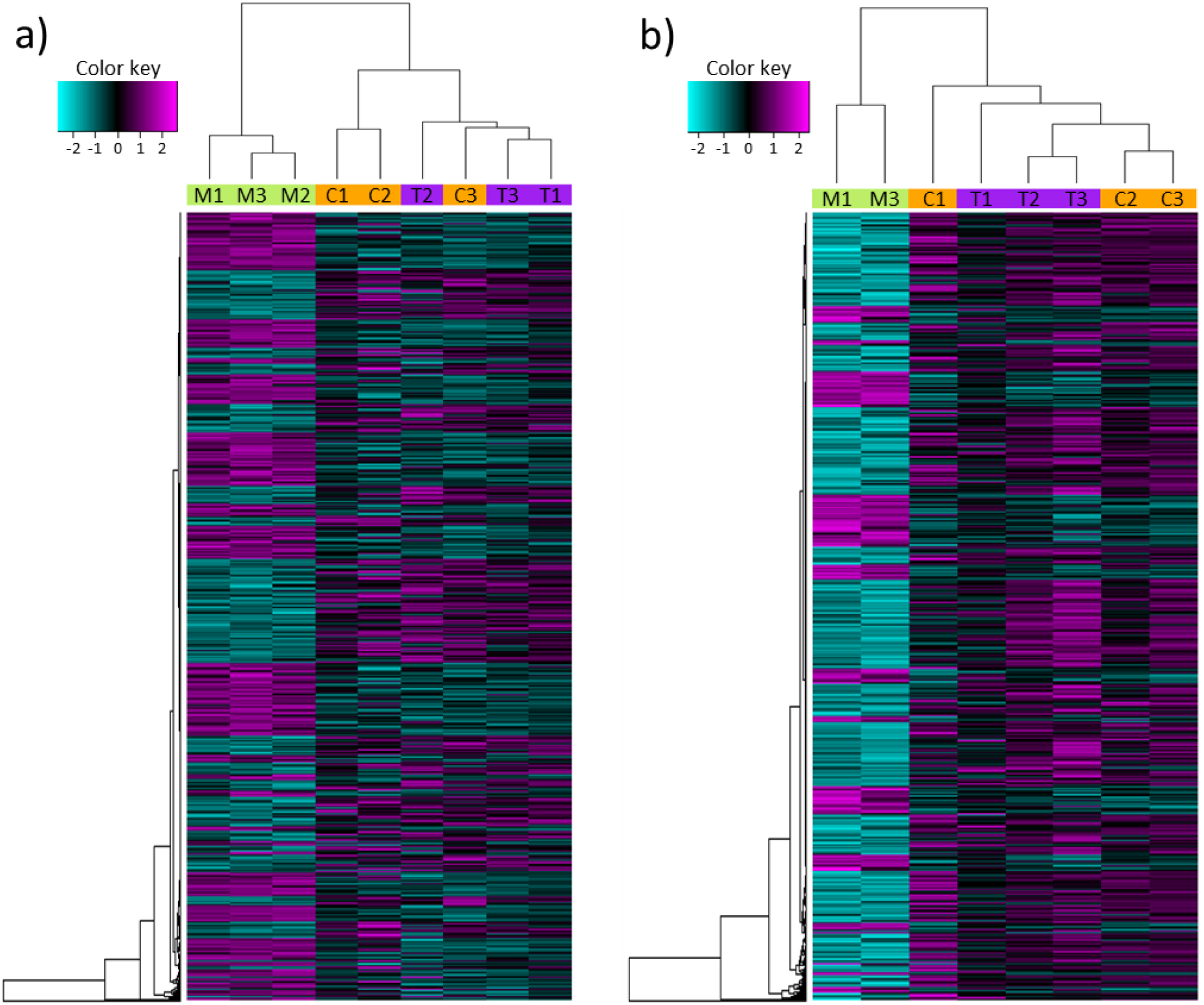
Hierarchical clustering of all differentially expressed genes (DEGs) in Myzus persicae feeding on CaMV- and TuYV-infected Arabidopsis thaliana (a) and Camelina sativa (b), compared to mock-inoculated control plants (Mock-inoculated [M], TuYV-infected [T] and CaMV-infected [C]) (Supplementary Dataset S1). The color key scale displays the row Z-score (normalized counts) from −2 to +2 as a gradient from cyan to magenta.

In summary, both GO and a general heatmap analyses indicated an important effect of plant infection on the aphid transcriptome, which was strongly shaped by the host plant identity (2/3 of the DEGs) and less so by the virus species (1/3 of the DEGs) and therefore the virus transmission mode.

### Discussion of DEGs by classes

In the following section, aphid DEGs were classified in several categories using as criteria whether or not the genes were differentially expressed in specific conditions (plant host species and virus identity) (Figure 4). The rationale was to identify genes that were general players in plant-virus-aphid interactions (i.e. deregulated by both viruses and on both host plants; Figure 4a) and genes that were specifically deregulated either by one virus species or by one host species. Then, we extracted aphid DEGs related to one virus and conserved regardless of the host plant to highlight virus species/transmission mode-specific genes that were not sensitive to the host plant identity (Figure 4b). Finally, we compared TuYV vs CaMV effects on each host plant to reveal additional, host plant-specific, ‘manipulation strategies’ linked to the virus transmission mode (Figure 4c).

**Figure 4.**
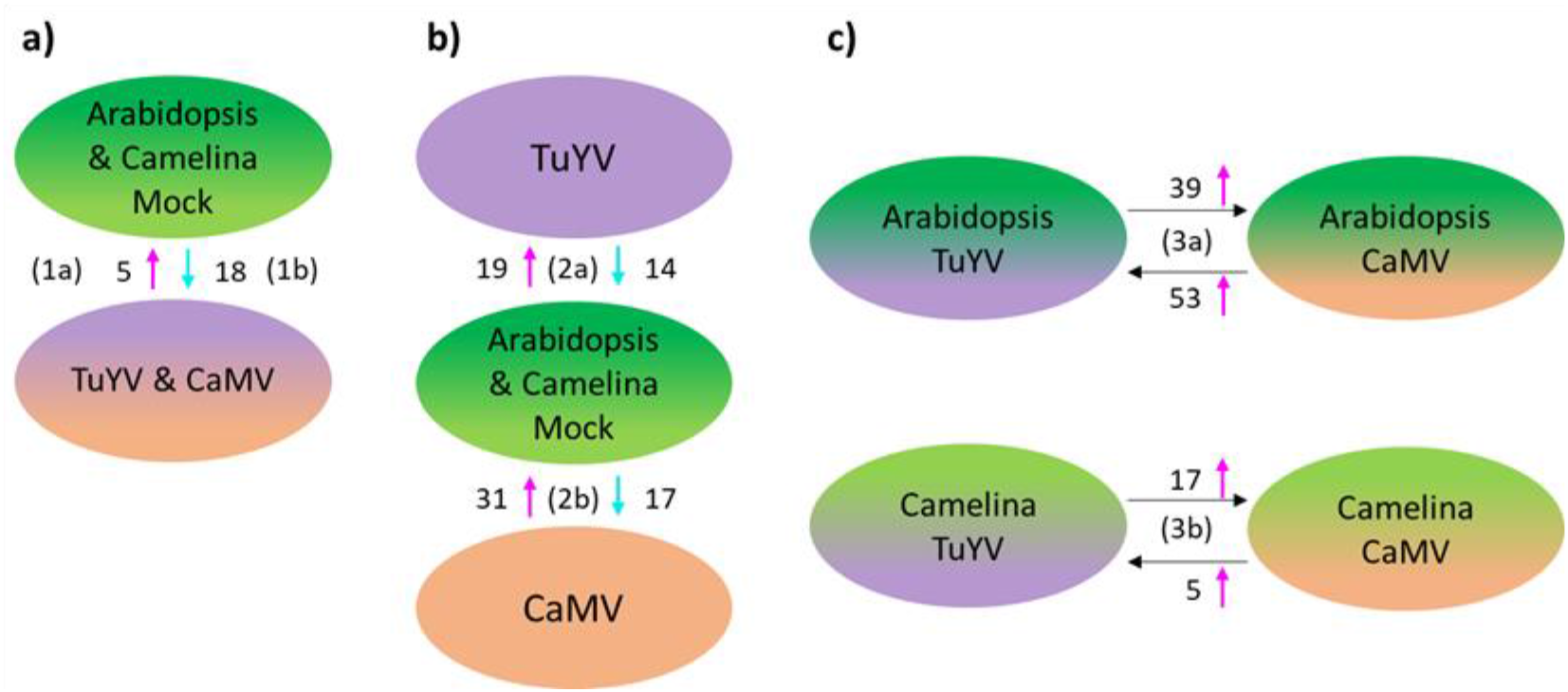
Summarizing figure presenting the number of differentially expressed aphid genes (log2FC > 0.5 or <0.5) discussed in each subsection of the discussion (1a, 1b, 2a, 2b, 3a, 3b). a) differentially expressed aphid genes in common for all conditions, b) virus-specific aphid DEGs common on both host-plants, and c) host plant-specific aphid DEGs for TuYV vs CaMV. Magenta arrows indicate the number of upregulated genes and the cyan arrows the number of downregulated genes.

#### 1 Common deregulated genes in aphids feeding on CaMV- and TuYV-infected Arabidopsis and Camelina

This analysis was carried out on genes differentially up- or downregulated under all conditions. No homolog was identified for up-regulated genes. In the case of downregulated genes, we found some genes homologs where one homolog was downregulated for one virus and another one for the other virus (Table 1). For example, we identified two potentially secreted homologous cathepsin B-like proteases (g8486 for aphids infesting TuYV-infected plants and g24532 for aphids infesting CaMV-infected plants). These homologs were included in the analysis. The rationale was that one specific host or infection condition might deregulate a specific gene but that the overall effect on plant aphid interactions might be the same or very similar for both genes (in this case the two cathepsin Bs might have a similar role as saliva effectors).

**Table 1.**
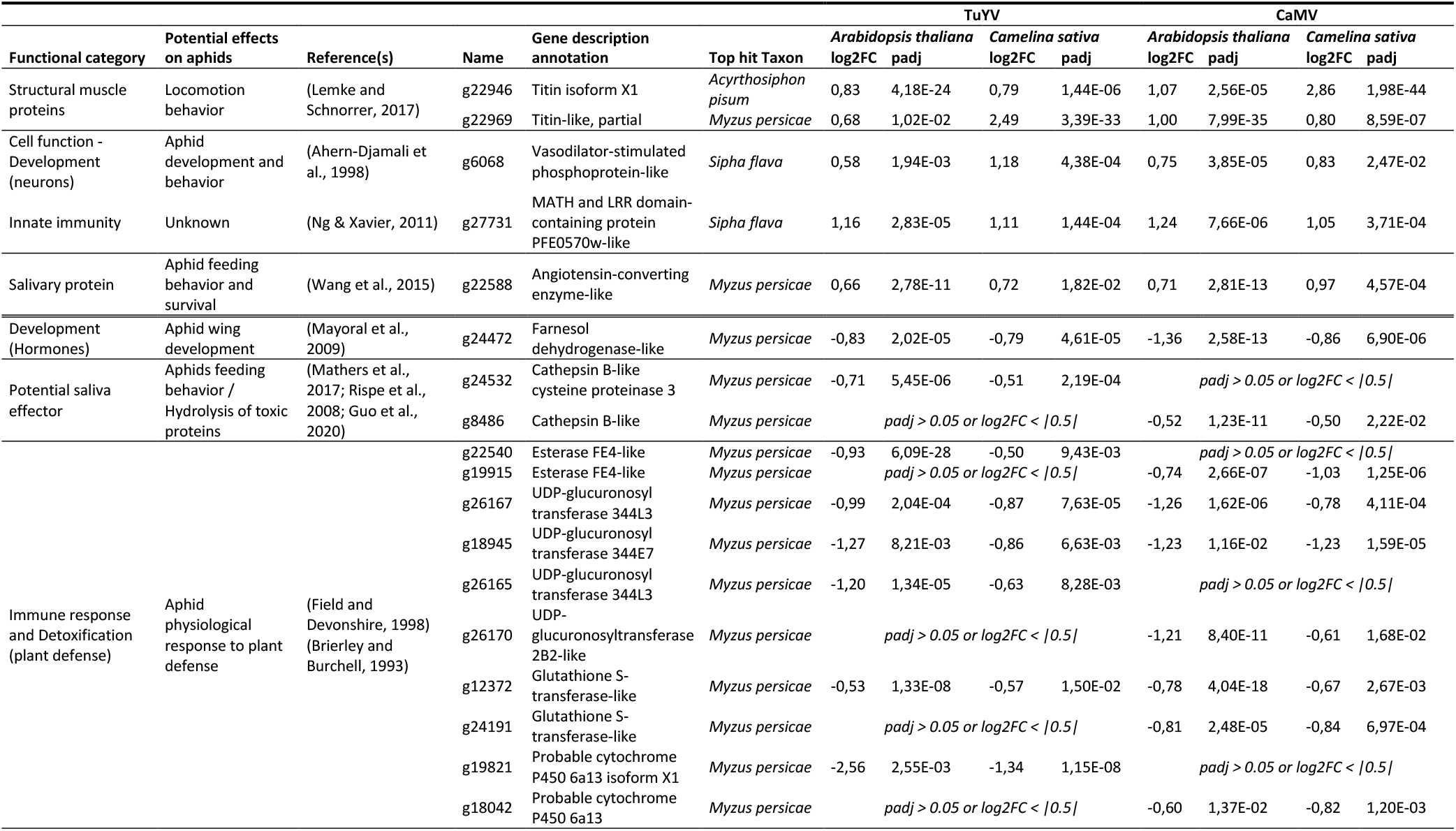
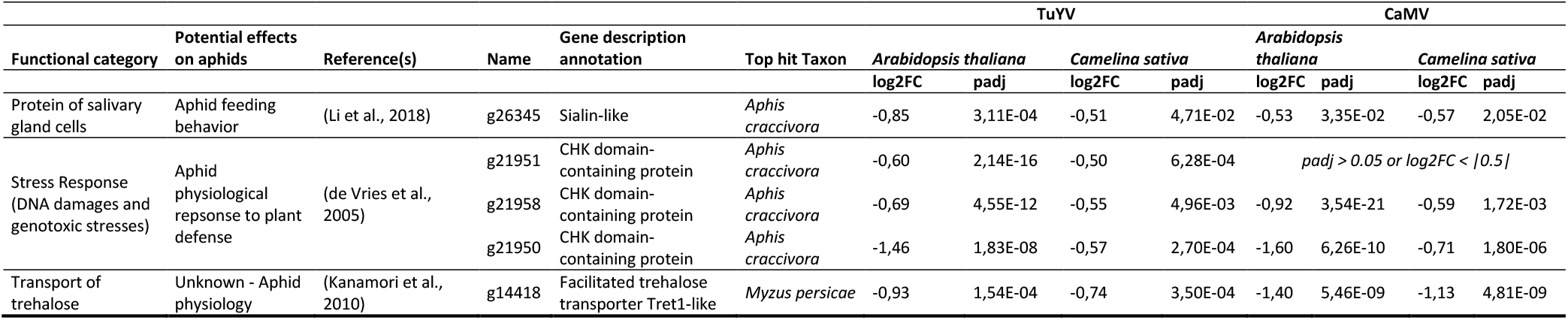
Selected differentially expressed aphid genes in common for aphids feeding on both CaMV and TuYV-infected Arabidopsis and Camelina. Single lines separate genes by functional categories. The double line separates up-regulated from down-regulated genes.

| Functional category | Potential effects on aphids | Reference(s) | Name | Gene description annotation | Top hit Taxon | TuYV |  |  |  | CaMV |  |  |  |
| --- | --- | --- | --- | --- | --- | --- | --- | --- | --- | --- | --- | --- | --- |
|  |  |  |  |  |  | Arabidopsis thaliana |  | Camelina sativa |  | Arabidopsis thaliana |  | Camelina sativa |  |
|  |  |  |  |  |  | log2FC | padj | log2FC | padj | log2FC | padj | log2FC | padj |
| Structural muscle proteins | Locomotion behavior | (Lemke and Schnorrer, 2017) | g22946 | Titin isoform X1 | *Acyrtosiphon pisum* | 0,83 | 4,18E-24 | 0,79 | 1,44E-06 | 1,07 | 2,56E-05 | 2,86 | 1,98E-44 |
|  |  |  | g22969 | Titin-like, partial | *Myzus persicae* | 0,68 | 1,02E-02 | 2,49 | 3,39E-33 | 1,00 | 7,99E-35 | 0,80 | 8,59E-07 |
| Cell function - Development (neurons) | Aphid development and behavior | (Ahern-Djamali et al., 1998) | g6068 | Vasodilator-stimulated phosphoprotein-like | *Sipha flava* | 0,58 | 1,94E-03 | 1,18 | 4,38E-04 | 0,75 | 3,85E-05 | 0,83 | 2,47E-02 |
| Innate immunity | Unknown | (Ng & Xavier, 2011) | g27731 | MATH and LRR domain-containing protein PFE0570w-like | *Sipha flava* | 1,16 | 2,83E-05 | 1,11 | 1,44E-04 | 1,24 | 7,66E-06 | 1,05 | 3,71E-04 |
| Salivary protein | Aphid feeding behavior and survival | (Wang et al., 2015) | g22588 | Angiotensin-converting enzyme-like | *Myzus persicae* | 0,66 | 2,78E-11 | 0,72 | 1,82E-02 | 0,71 | 2,81E-13 | 0,97 | 4,57E-04 |
| Development (Hormones) | Aphid wing development | (Mayoral et al., 2009) | g24472 | Farnesol dehydrogenase-like | *Myzus persicae* | -0,83 | 2,02E-05 | -0,79 | 4,61E-05 | -1,36 | 2,58E-13 | -0,86 | 6,90E-06 |
| Potential saliva effector | Aphids feeding behavior / Hydrolysis of toxic proteins | (Mathers et al., 2017; Rispe et al., 2008; Guo et al., 2020) | g24532 | Cathepsin B-like cysteine proteinase 3 | *Myzus persicae* | -0,71 | 5,45E-06 | -0,51 | 2,19E-04 | *padj > 0.05 or log2FC < \|0.5\|* |  |  |  |
|  |  |  | g8486 | Cathepsin B-like | *Myzus persicae* | *padj > 0.05 or log2FC < \|0.5\|* |  |  |  | -0,52 | 1,23E-11 | -0,50 | 2,22E-02 |
| Immune response and Detoxification (plant defense) | Aphid physiological response to plant defense | (Field and Devonshire, 1998) (Brierley and Burchell, 1993) | g22540 | Esterase FE4-like | *Myzus persicae* | -0,93 | 6,09E-28 | -0,50 | 9,43E-03 | *padj > 0.05 or log2FC < \|0.5\|* |  |  |  |
|  |  |  | g19915 | Esterase FE4-like | *Myzus persicae* | *padj > 0.05 or log2FC < \|0.5\|* |  |  |  | -0,74 | 2,66E-07 | -1,03 | 1,25E-06 |
|  |  |  | g26167 | UDP-glucuronosyl transferase 344L3 | *Myzus persicae* | -0,99 | 2,04E-04 | -0,87 | 7,63E-05 | -1,26 | 1,62E-06 | -0,78 | 4,11E-04 |
|  |  |  | g18945 | UDP-glucuronosyl transferase 344E7 | *Myzus persicae* | -1,27 | 8,21E-03 | -0,86 | 6,63E-03 | -1,23 | 1,16E-02 | -1,23 | 1,59E-05 |
|  |  |  | g26165 | UDP-glucuronosyl transferase 344L3 | *Myzus persicae* | -1,20 | 1,34E-05 | -0,63 | 8,28E-03 | *padj > 0.05 or log2FC < \|0.5\|* |  |  |  |
|  |  |  | g26170 | UDP-glucuronosyltransferase 2B2-like | *Myzus persicae* | *padj > 0.05 or log2FC < \|0.5\|* |  |  |  | -1,21 | 8,40E-11 | -0,61 | 1,68E-02 |
|  |  |  | g12372 | Glutathione S-transferase-like | *Myzus persicae* | -0,53 | 1,33E-08 | -0,57 | 1,50E-02 | -0,78 | 4,04E-18 | -0,67 | 2,67E-03 |
|  |  |  | g24191 | Glutathione S-transferase-like | *Myzus persicae* | *padj > 0.05 or log2FC < \|0.5\|* |  |  |  | -0,81 | 2,48E-05 | -0,84 | 6,97E-04 |
|  |  |  | g19821 | Probable cytochrome P450 6a13 isoform X1 | *Myzus persicae* | -2,56 | 2,55E-03 | -1,34 | 1,15E-08 | *padj > 0.05 or log2FC < \|0.5\|* |  |  |  |
|  |  |  | g18042 | Probable cytochrome P450 6a13 | *Myzus persicae* | *padj > 0.05 or log2FC < \|0.5\|* |  |  |  | -0,60 | 1,37E-02 | -0,82 | 1,20E-03 |
|  |  |  |  |  |  | *Arabidopsis thaliana* |  | *Camelina sativa* |  | *Arabidopsis thaliana* |  | *Camelina sativa* |  |
|  |  |  |  |  |  | log2FC | padj | log2FC | padj | log2FC | padj | log2FC | padj |
| Protein of salivary gland cells | Aphid feeding behavior | (Li et al., 2018) | g26345 | Sialin-like | *Aphis craccivora* | -0,85 | 3,11E-04 | -0,51 | 4,71E-02 | -0,53 | 3,35E-02 | -0,57 | 2,05E-02 |
| Stress Response (DNA damages and genotoxic stresses) | Aphid physiological response to plant defense | (de Vries et al., 2005) | g21951 | CHK domain-containing protein | *Aphis craccivora* | -0,60 | 2,14E-16 | -0,50 | 6,28E-04 | *padj > 0.05 or log2FC < \|0.5\|* |  |  |  |
|  |  |  | g21958 | CHK domain-containing protein | *Aphis craccivora* | -0,69 | 4,55E-12 | -0,55 | 4,96E-03 | -0,92 | 3,54E-21 | -0,59 | 1,72E-03 |
|  |  |  | g21950 | CHK domain-containing protein | *Aphis craccivora* | -1,46 | 1,83E-08 | -0,57 | 2,70E-04 | -1,60 | 6,26E-10 | -0,71 | 1,80E-06 |
| Transport of trehalose | Unknown - Aphid physiology | (Kanamori et al., 2010) | g14418 | Facilitated trehalose transporter Tret1-like | *Myzus persicae* | -0,93 | 1,54E-04 | -0,74 | 3,50E-04 | -1,40 | 5,46E-09 | -1,13 | 4,81E-09 |

##### 1a Aphid genes UPREGULATED by both VIRUSES on both PLANTS

Only five genes were upregulated in Myzus feeding on both CaMV- and TuYV-infected vs mock-inoculated plants. Two of them (g22946 and g22969) code for titins, which are structural muscle proteins (Lemke & Schnorrer, 2017) (Table 1). Their upregulation could potentially affect locomotion behavior and facilitate intra or inter-plants vector movement, as was recently observed for TuYV-viruliferous aphids (Chesnais et al., 2020). The third commonly upregulated gene (g6068) codes for a vasodilator-stimulated phosphoprotein-like (VASP) protein, which is associated with actin filaments and focal adhesions (Ahern-Djamali et al., 1998). It can participate in neural development and function, as suggested for its Drosophila homolog Ena (Ahern-Djamali et al., 1998). Transferred to the present work, VASP might be induced by viruses to modulate aphid development and behavior, a feature that has been reported several times in the recent literature (Mauck et al., 2018). The fourth gene upregulated in all modalities encodes an angiotensin-converting enzyme-like protein (g22588), the orthologue of *Acyrthosiphon pisum* ACE1. g22588 contains a signal peptide and could therefore be a saliva protein. *Acyrthosiphon pisum* ACE1 is expressed in salivary glands and modulates aphid-plant interactions by affecting the feeding behavior and survival of aphids on host plants (Wang, Luo, et al., 2015). More precisely, silencing of ACE1 and ACE2 shortened the lifespan of *A. pisum* on plants but not in membrane feeding assays. Thus, the ACE1 g22588 possibly counteracts plant defenses and its overexpression could help Myzus to better cope with plant defenses and to increase its lifespan. ACE and other metalloproteases have also been found in the saliva of other phytophagous and blood-feeding arthropods (Decrem et al., 2008; Stafford-Banks et al., 2014) and are believed to be part of their arsenal counteracting defense responses of their hosts (reviewed by Hopp & Sinnis, 2015; Wang et al., 2017; Chen & Mao, 2020; Pham et al., 2021). Thus, the increased expression of ACE, as observed here in the interaction of Myzus with two viruses and on two host plants, could be a common ‘manipulation strategy’ shared among plant viruses to facilitate aphid feeding on virus-infected host plants and accelerate virus acquisition. The fifth commonly upregulated gene is the uncharacterized Myzus gene g27731 encoding a protein with MATH and LRR domains. Since the LRR domain is evolutionary conserved in many proteins associated with innate immunity pathways (Ng & Xavier, 2011), this protein might be a good candidate for further studies on virus-mediated manipulation of insect vectors.

##### 1b Aphid genes DOWNREGULATED by both VIRUSES on both PLANTS

We found 18 common downregulated genes (including some homologs) in aphids fed on virus-infected plants, 13 of which are implicated in aphids’ physiological responses to plant defenses, i.e. stress-related genes (Table 1). Previously, we have demonstrated that plant infection with CaMV and TuYV alters primary and secondary metabolism in Arabidopsis and Camelina strongly (Chesnais, Golyaev, et al., 2022). Consequently, aphids could also be stressed on the infected plants because of these important alterations of the plants. However, we found that stress-related aphid genes were downregulated in Myzus feeding on virus-infected plants. This suggests that plant infection with TuYV or CaMV could facilitate aphid infestation. The downregulated stress genes included those encoding the metabolic enzymes cytochrome P450s, glutathione-S-transferase and UDP-glucuronosyl transferases which can play a key role in detoxifying plant secondary metabolites (Brierley & Burchell, 1993; Li et al., 2007). We also noticed downregulation of genes encoding FE4-like esterases that belong to another class of enzymes involved in detoxification and that can confer insecticide resistance in Myzus (Field & Devonshire, 1998). Likewise, Myzus genes coding for CHK domain-containing proteins were downregulated (Table 1). CHKs (checkpoint kinases) are major mediators of cell cycle checkpoints in response to genotoxic and other stresses (de Vries et al., 2005).

Another aphid gene downregulated under all conditions encodes for a facilitated transmembrane trehalose transporter, Tret1. Trehalose (α-D-glucopyranosyl-(1,1)-α-D-glucopyranoside) is the main hemolymph sugar, and Tret1 is necessary for the transport of trehalose produced in the fat body and its uptake into other tissues that require a carbon source (Kanamori et al., 2010). Deregulation of this gene following virus acquisition has already been reported in other insect vectors and is not linked to the virus species or the transmission mode (e.g. Widana Gamage et al., 2018; Ding et al., 2019).

As mentioned above, we observed downregulation of distinct Cathepsin B3 (CathB)-encoding genes in aphids feeding on TuYV- or CaMV-infected vs mock-inoculated plants. CathBs are detoxifying proteases found in saliva and intestine and are subject to gene amplification, which is thought to be an adaptation of saliva and intestine of aphids feeding on phloem sap from different plant species (Rispe et al., 2007; Mathers et al., 2017). The CathB identified in Myzus feeding on TuYV-infected plants (g24532) is the same as the one described by Guo et al. (2020), and the one found in Myzus infesting CaMV-infected plants (g8486) is closely related to it (87 % identity on the amino acid level). The latter paralog (CathB3) is a saliva protease and an effector that induces plant defenses. Therefore, its downregulation can be proviral by facilitating plant infestation. Up- or downregulation of CathB-encoding genes during host-virus interactions has been observed in many arthropod vectors (aphids, whitefly, thrips, leafhoppers, mites and mosquitos), suggesting importance of the cathepsin Bs in virus-host-vector interactions and, possibly, transmission (Pinheiro et al., 2017; Hasegawa et al., 2018; Widana Gamage et al., 2018; Gupta et al., 2019; Caicedo et al., 2019; Li et al., 2019, 2020; Xu et al., 2021). Another feeding-related downregulated gene in aphids infesting virus-infected Arabidopsis and Camelina encodes a sialin (g26345). A mammalian ortholog of the sialin gene encodes a membrane protein of salivary gland cells controlling osmolarity and composition of saliva (Li et al., 2018). Finally, among commonly downregulated Myzus genes we identified a gene encoding a farnesol dehydrogenase-like protein (g24472), implicated in hormone metabolism. This gene could be responsible for the oxidation of farnesol to farnesal, a precursor of the juvenile hormone as shown for mosquitoes (Mayoral et al., 2009). Downregulation of juvenile hormone can favor wing development (Zhang et al., 2019), which might facilitate viral spread.

Taken together, we observed that among the ‘common’ genes those involved in locomotion, neural development and lifespan were rather upregulated in aphids feeding on virus-infected plants. This might favor aphid mobility and survival and in turn virus dispersion. Genes involved in stress responses and saliva functions were mostly downregulated (except the saliva protein ACE1 contributing to lifespan), indicating that viral infection facilitates aphid infestation of the host plants, for example by dampening anti-herbivore plant defenses as observed in our previous study (Chesnais, Golyaev, et al., 2022).

#### 2 Virus-specific aphid DEGs on both host-plants

##### 2a TuYV-specific DEGs in Myzus feeding on Arabidopsis and Camelina

To know whether plant viruses can impact aphid genes independently of the plant host, we first screened for aphid DEGs in common for aphids feeding on TuYV-infected Camelina and Arabidopsis. We found 19 upregulated genes (see a complete list in Table S3). Two of them might influence aphid feeding behavior (Table 2a). One (g26473) codes for a putative stylet sheath protein. Stylet sheaths are formed by gelling saliva that is secreted during stylet penetration in plant tissue. The sheaths insulate the stylets and potentially protect them from plant defenses and seal cell and phloem puncture sites (Will et al., 2012). Silencing of an *A. pisum* sheath protein gene disrupted sheath formation and disturbed phloem-feeding (Will & Vilcinskas, 2015), suggesting that upregulation, as observed here, might conversely facilitate and accelerate aphid feeding behavior on TuYV-infected plants (as observed by Chesnais et al., 2020), and hence TuYV acquisition. The second TuYV-specific feeding-related aphid gene (g15241) codes for a receptor for the insect neuropeptide SIFamide that might control feeding indirectly by modulating behavior, as shown for SIFamide in the Chagas disease vector, the kissing bug *Rhodnius prolixus* (Ayub et al., 2020). The other upregulated genes are mostly related to development. Interestingly, two of them, forkhead box protein O (Foxo) and ATP-binding cassette sub-family G member 4-like (ABCG4) could be involved in aphid wing formation (Shang et al., 2020; Grantham et al., 2020). Induction of wings could considerably increase virus propagation by aphids, especially over long distances, as recently shown for CMV transmission (Jayasinghe et al., 2021). In this specific case, the wing formation was attributed to a virus satellite co-infecting the plant. The few downregulated genes (n = 14, see a complete list in Table S3) specific for aphids on TuYV-infected plants are involved in detoxification and are closely related to the detoxification genes downregulated by both viruses in all conditions (see the previous section).

**Table 2.** Selected genes commonly deregulated in aphids feeding on a) TuYV-infected and b) CaMV-infected host plants (Arabidopsis and Camelina). Single lines separate genes by functional categories.

| a) TuYV |  |  |  |  |  | Arabidopsis thaliana |  | Camelina sativa |  |
| --- | --- | --- | --- | --- | --- | --- | --- | --- | --- |
| Functional category | Potential effects on aphids | Reference(s) | Name | Gene description annotation | Top hit Taxon | log2FC | padj | log2FC | padj |
| Saliva protein | Aphid feeding behavior | (Will et al., 2012) | g26473 | Putative sheath protein, partial | *Sitobion avenae* | 0,52 | 1,76E-11 | 0,69 | 9,93E-03 |
| Insect neuropeptide | Aphid behavior (sleep, sexual and feeding) | (Ayub et al., 2020) | g15241 | Neuropeptide SIFamide receptor-like | *Myzus persicae* | 0,57 | 5,75E-03 | 1,22 | 4,50E-03 |
| Membrane-associated transporter | Aphid development (wing) | (Shang et al., 2020; Jayasinghe et al., 2021) | g16568 | ATP-binding cassette sub-family G member 4-like | *Myzus persicae* | 0,52 | 2,15E-15 | 0,82 | 2,76E-07 |
| Transcription factor | Aphid development (wing) | (Grantham et al., 2020) | g24925 | Forkhead box protein O | *Myzus persicae* | 0,50 | 9,81E-04 | 1,11 | 3,62E-06 |
| b) CaMV |  |  |  |  |  | Arabidopsis thaliana |  | Camelina sativa |  |
| Functional category | Potential effects on aphids | Reference(s) | Name | Gene description annotation | Top hit Taxon | log2FC | padj | log2FC | padj |
| Development (multiple pathways) | Aphid behavior and development |  | g19210 | Glucose dehydrogenase [FAD, quinone]-like isoform X1 | *Myzus persicae* | 0,87 | 6,77E-10 | 1,20 | 7,11E-04 |
|  |  |  | g19209 | Glucose dehydrogenase [FAD, quinone]-like | *Myzus persicae* | 0,51 | 6,21E-03 | 0,98 | 8,07E-04 |
| Structural protein | Aphid development - Virus interaction | (Deshoux et al., 2018) | g21498 | RR-2 cuticle protein 3, partial | *Myzus persicae* | 1,03 | 1,21E-10 | 0,85 | 9,89E-04 |
| Saliva protein | Aphid feeding behavior | (Huang et al., 2017; Shangguan et al., 2018) | g27683 | Mucin-2-like | *Myzus persicae* | 1,01 | 9,40E-42 | 1,54 | 9,71E-06 |
| Metalloproteases - Secreted protein | Aphid feeding behavior - Digestion | (Sterchi et al., 2008) | g7709 | Astacin-like | *Myzus persicae* | 1,13 | 1,26E-09 | 0,93 | 1,69E-02 |
| Saliva protein - Lipase activity | Aphid feeding behavior | (Chaudhary et al., 2015) | g16515 | Pancreatic lipase-related protein 2-like | *Myzus persicae* | -0,71 | 8,11E-14 | -0,50 | 8,80E-03 |
| Saliva protein | Aphid feeding behavior | (Enders et al., 2015; Champagne et al., 1995) | g22531 | Protein 5NUC isoform X1 | *Acyrtosiphon pisum* | -0,62 | 4,17E-02 | -0,70 | 6,29E-04 |
| Carbohydrate metabolism | Aphid metabolism | (Qin et al., 2018) | g20667 | Sugar transporter SWEET1-like | *Myzus persicae* | -0,53 | 1,87E-08 | -0,54 | 1,18E-02 |

##### 2b CaMV-specific DEGs in Myzus feeding on Arabidopsis and Camelina

We also analyzed the common and specific DEGs only found in aphids fed on CaMV-infected Arabidopsis and Camelina. We identified a total of 48 DEGs (31 upregulated and 17 downregulated genes, see the complete list in Table S4). One of the upregulated genes codes for a glucose dehydrogenase (Table 2b). Since glucose dehydrogenases are involved in multiple pathways, it is difficult to attribute a precise role for these enzymes in CaMV-aphid interactions. Several other upregulated genes might modulate aphid development and feeding behavior. For example, the gene g21498 codes for a structural RR-2 cuticle protein 3. Other RR-1 and RR-2 cuticle proteins are involved in virus–vector interactions (Deshoux et al., 2018), and it would be interesting to investigate a possible role of the RR-2 cuticle protein 3 in CaMV transmission. Another upregulated gene codes for an astacin (g7709), which belongs to a group of metalloproteases with various functions (Sterchi et al., 2008). Since the astacin identified here contains a signal peptide for secretion, it is tempting to speculate that it could be released during salivation or digestion and that upregulation might improve feeding.

As mentioned in the above section, both TuYV and CaMV infections deregulated some aphid genes linked to salivary proteins (for example ACE1 and CathB). CaMV acquisition, but not TuYV acquisition, upregulated in Myzus another potential saliva gene coding for a mucin-2-like protein (g27683). In animals, saliva mucins protect by lubrication soft and hard tissues in the mouth (Turner, 2016). Their aphid homologs could have similar functions in protecting the stylet surface. Insect mucins have been studied thoroughly in the brown planthopper *Nilaparvata lugens*. One of them, NlMul, is a major component of watery and gelling saliva, required for proper feeding (Huang et al., 2017). Another one, NlMLP, is also involved in sheath formation, but in addition this mucin elicits plant defense responses (Shangguan et al., 2018). Transferred to aphids, genes encoding mucins could be involved in the significant phagostimulation observed in aphids on CaMV-infected plants compared to healthy plants (Chesnais et al., 2021).

Among genes downregulated by CaMV (but not by TuYV) in Myzus when feeding on both hosts were other genes coding for potential saliva proteins. One of them (g22531) codes for a 5’-nucleotidase with some similarities to a 5’-nucleotidase downregulated by various stresses in *A. glycines* (Enders et al., 2015) and to a saliva-contained 5-’nucleotidase of the mosquito *Aedes aegypti* (Champagne et al., 1995). A second gene codes for a pancreatic lipase-related 2-like protein (g16515). Similar enzymes have been identified in the salivary proteome of the potato aphid *Macrosiphum euphorbiae* (Chaudhary et al., 2015). Other pancreatic lipases are involved in vector interactions with circulative viruses. A pancreatic lipase from *Rhopalosiphum padi* binds to the CP and RT of barley yellow dwarf virus (family *Luteoviridae*) in yeast two-hybrid assay (Wang, Wu, et al., 2015) and the gene expression of another pancreatic lipase is downregulated in *Bemisia tabaci* fed on TYLCV-infected tomato (Hasegawa et al., 2018). Thus, an impact of downregulation of this gene on non-circulative CaMV transmission could be indirect. Among other downregulated genes was the sugar transporter SWEET1-like gene (g20667), which codes for the midgut receptor of at least three planthopper-transmitted circulative, propagative viruses (Qin et al., 2018). A role, if any, for this gene in non-circulative transmission of CaMV by aphids could also be indirect, possibly by increasing feeding activity and concomitant virus acquisition, due to reduced sugar uptake.

#### 3 Host plant-specific aphid DEGs for TuYV vs CaMV

To reveal an additional, host plant-specific, contribution to viral manipulation strategies linked to circulative vs non-circulative transmission modes, we analyzed DEGs in aphids feeding on TuYV- vs CaMV-infected Arabidopsis and in aphids feeding on TuYV- vs CaMV-infected Camelina (Figure 1e, TuYV vs CaMV). Since for Arabidopsis the total number of such aphid DEGs was 380, we applied a cut-off of log2FC (fold changes) > 0.5 for upregulated genes and < −0.5 for downregulated genes to limit the number to 90 genes. This step was not necessary in the case of Camelina, where in total only 22 aphid DEGs were observed (see the complete lists in Tables S3, S4 and S5).

##### 3a TuYV vs CaMV in Arabidopsis

A higher proportion of genes was upregulated in aphids feeding on TuYV-infected Arabidopsis, compared to aphids infesting CaMV-infected Arabidopsis (see Table 3a and Supplementary Table S5-6). Two of them (g5369 and g10419) encode chitinases that are essential for insect survival, molting and development (Arakane & Muthukrishnan, 2010). Four other genes encode the development-related proteins octopamine receptor Oamb (g15146), homeotic protein distal-less-like protein (g5303), zinc finger protein Elbow (g24564) and bombyxin C-2 like protein (g7214) (Campbell & Tomlinson, 1998; Weihe et al., 2004; Wang et al., 2016; Ding et al., 2017). Thus, compared to CaMV, TuYV infection of Arabidopsis specifically induces higher expression of aphid genes potentially involved in wing formation/development. This could promote, as discussed above, the formation of alate individuals with consequences on TuYV dispersal to new plants.

**Table 3.**
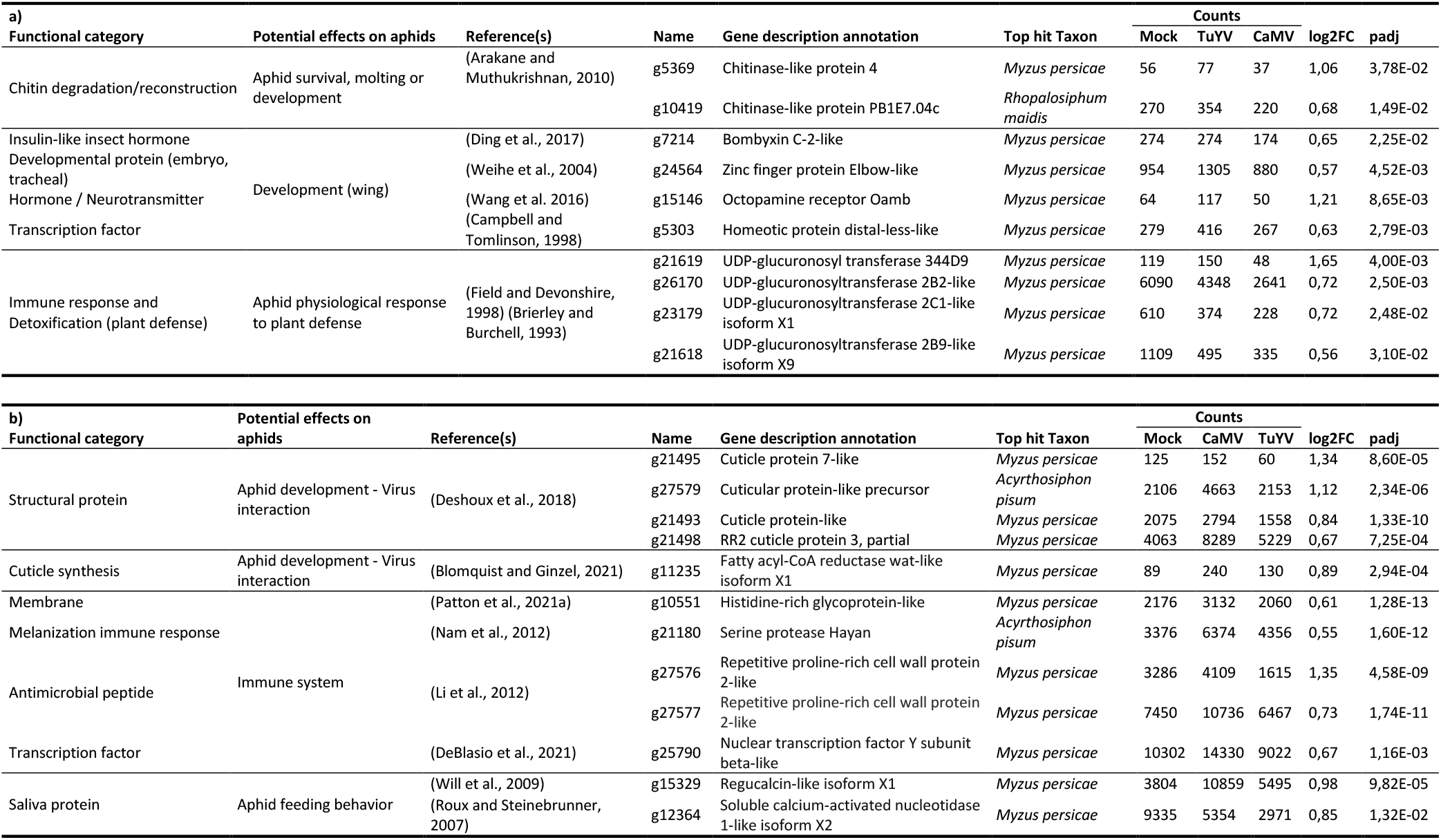
Selected genes differentially expressed in aphids feeding on TuYV-infected vs CaMV-infected Arabidopsis. a) up-regulated on TuYV-infected Arabidopsis and b) up-regulated on CaMV-infected Arabidopsis. Single lines separate genes by functional categories.

| a) |  |  |  |  |  | Counts |  |  |  |  |
| --- | --- | --- | --- | --- | --- | --- | --- | --- | --- | --- |
| Functional category | Potential effects on aphids | Reference(s) | Name | Gene description annotation | Top hit Taxon | Mock | TuYV | CaMV | log2FC | padj |
| Chitin degradation/reconstruction | Aphid survival, molting or development | (Arakane and Muthukrishnan, 2010) | g5369 | Chitinase-like protein 4 | *Myzus persicae* | 56 | 77 | 37 | 1,06 | 3,78E-02 |
|  |  |  | g10419 | Chitinase-like protein PB1E7.04c | *Rhopalosiphum maidis* | 270 | 354 | 220 | 0,68 | 1,49E-02 |
| Insulin-like insect hormone | Development (wing) | (Ding et al., 2017) | g7214 | Bombyxin C-2-like | *Myzus persicae* | 274 | 274 | 174 | 0,65 | 2,25E-02 |
| Developmental protein (embryo, tracheal) |  | (Weihe et al., 2004) | g24564 | Zinc finger protein Elbow-like | *Myzus persicae* | 954 | 1305 | 880 | 0,57 | 4,52E-03 |
| Hormone / Neurotransmitter |  | (Wang et al. 2016) | g15146 | Octopamine receptor Oamb | *Myzus persicae* | 64 | 117 | 50 | 1,21 | 8,65E-03 |
| Transcription factor |  | (Campbell and Tomlinson, 1998) | g5303 | Homeotic protein distal-less-like | *Myzus persicae* | 279 | 416 | 267 | 0,63 | 2,79E-03 |
| Immune response and Detoxification (plant defense) | Aphid physiological response to plant defense | (Field and Devonshire, 1998) (Brierley and Burchell, 1993) | g21619 | UDP-glucuronosyl transferase 344D9 | *Myzus persicae* | 119 | 150 | 48 | 1,65 | 4,00E-03 |
|  |  |  | g26170 | UDP-glucuronosyltransferase 2B2-like | *Myzus persicae* | 6090 | 4348 | 2641 | 0,72 | 2,50E-03 |
|  |  |  | g23179 | UDP-glucuronosyltransferase 2C1-like isoform X1 | *Myzus persicae* | 610 | 374 | 228 | 0,72 | 2,48E-02 |
|  |  |  | g21618 | UDP-glucuronosyltransferase 2B9-like isoform X9 | *Myzus persicae* | 1109 | 495 | 335 | 0,56 | 3,10E-02 |
| b) |  |  |  |  |  | Counts |  |  |  |  |
| Functional category | Potential effects on aphids | Reference(s) | Name | Gene description annotation | Top hit Taxon | Mock | CaMV | TuYV | log2FC | padj |
| Structural protein | Aphid development - Virus interaction | (Deshoux et al., 2018) | g21495 | Cuticle protein 7-like | *Myzus persicae* | 125 | 152 | 60 | 1,34 | 8,60E-05 |
|  |  |  | g27579 | Cuticular protein-like precursor | *Acyrtosiphon pisum* | 2106 | 4663 | 2153 | 1,12 | 2,34E-06 |
|  |  |  | g21493 | Cuticle protein-like | *Myzus persicae* | 2075 | 2794 | 1558 | 0,84 | 1,33E-10 |
|  |  |  | g21498 | RR2 cuticle protein 3, partial | *Myzus persicae* | 4063 | 8289 | 5229 | 0,67 | 7,25E-04 |
| Cuticle synthesis | Aphid development - Virus interaction | (Blomquist and Ginzel, 2021) | g11235 | Fatty acyl-CoA reductase wat-like isoform X1 | *Myzus persicae* | 89 | 240 | 130 | 0,89 | 2,94E-04 |
| Membrane | Immune system | (Patton et al., 2021a) | g10551 | Histidine-rich glycoprotein-like | *Myzus persicae* | 2176 | 3132 | 2060 | 0,61 | 1,28E-13 |
| Melanization immune response |  | (Nam et al., 2012) | g21180 | Serine protease Hayan | *Acyrtosiphon pisum* | 3376 | 6374 | 4356 | 0,55 | 1,60E-12 |
| Antimicrobial peptide |  | (Li et al., 2012) | g27576 | Repetitive proline-rich cell wall protein 2-like | *Myzus persicae* | 3286 | 4109 | 1615 | 1,35 | 4,58E-09 |
|  |  |  | g27577 | Repetitive proline-rich cell wall protein 2-like | *Myzus persicae* | 7450 | 10736 | 6467 | 0,73 | 1,74E-11 |
| Transcription factor |  | (DeBlasio et al., 2021) | g25790 | Nuclear transcription factor Y subunit beta-like | *Myzus persicae* | 10302 | 14330 | 9022 | 0,67 | 1,16E-03 |
| Saliva protein | Aphid feeding behavior | (Will et al., 2009) | g15329 | Regucalcin-like isoform X1 | *Myzus persicae* | 3804 | 10859 | 5495 | 0,98 | 9,82E-05 |
|  |  | (Roux and Steinebrunner, 2007) | g12364 | Soluble calcium-activated nucleotidase 1-like isoform X2 | *Myzus persicae* | 9335 | 5354 | 2971 | 0,85 | 1,32E-02 |

Interestingly, Myzus feeding on CaMV-infected Arabidopsis showed a different subset of developmental genes expressed at higher levels than Myzus feeding on TuYV-infected Arabidopsis. Four of these genes encode cuticle proteins (Table 3b and Supplementary Table S6). The fatty acyl-CoA reductase wat-like isoform X1 gene (g11235) that was also expressed at a higher level in the presence of CaMV, compared to TuYV, belongs to a gene family mediating the synthesis of insect cuticular hydrocarbons that are involved in the waterproofing of insect cuticles but also functions in signaling (Blomquist & Ginzel, 2021).

In addition to developmental genes, nine Myzus genes related to defense and detoxification responses were differentially expressed in Myzus after acquisition of TuYV on Arabidopsis, compared to acquisition of CaMV on Arabidopsis (Table 3a,b and Supplementary Table S5). For example, variable deregulations in the different conditions were observed for four genes of the UDP-glucuronosyl transferase gene family encoding detoxification enzymes (Brierley & Burchell, 1993). However, other genes that could be related to defense and/or detoxification were expressed at higher levels in CaMV-exposed aphids compared to aphids fed on TuYV-infected plants, such as the gene encoding the Hayan serine protease (g21180), which activates the melanization immune response to physical or septic wounding (Nam et al., 2012) and a gene encoding a histidine-rich glycoprotein (g10551). A gene coding for an anti-microbial peptide, repetitive proline-rich cell wall protein 2-like (g27577) (Li et al., 2012), was also expressed at higher levels in aphids fed on CaMV-infected plants than in aphids fed on TuYV-infected plants. A similar trend was found for the nuclear transcription factor Y subunit beta-like (g25790), which might interact with PLRV virions (DeBlasio et al., 2021) and a homolog of which belongs to the upregulated genes associated with the KEGG category “viral infectious disease” in whiteflies feeding on tomato infected with semi-persistent cucurbit yellow stunting disorder virus (genus *Crinivirus*, family *Closteroviridae*) (Kaur et al., 2019). Overall, we observed that different immune defense and detoxification pathways are affected in Myzus feeding on CaMV-infected Arabidopsis, compared to Myzus feeding on TuYV-infected Arabidopsis. This might be related to the different transmission modes of the two viruses. TuYV being circulative is expected to interact intimately with the vector and maybe even evade immune responses. On the other hand, CaMV interaction with the vector is confined to the stylet tip. Therefore, CaMV might rather modulate feeding responses. This might be illustrated by the strong activation of saliva genes (see below) following CaMV acquisition, whereas the impact of CaMV on developmental genes was comparably low. However, one needs to keep in mind that we discuss here only a subset of 90 most strongly deregulated genes in CaMV- exposed aphids compared to TuYV-exposed aphids.

Interestingly, in aphids feeding on CaMV-infected Arabidopsis, considerably more genes related to salivary proteins were expressed at higher levels, compared to those feeding on TuYV-infected Arabidopsis. Salivary proteins, liberated in the apoplast and plant cells or in the phloem during aphid probing and feeding activity, respectively, are excellent candidates to target defense pathways directly in the plant. Among them was the gene encoding a regucalcin (g15329) that has been identified earlier in the saliva of other aphid species (van Bel & Will, 2016). Regucalcin and other calcium-binding proteins could reduce calcium availability in the phloem, and subsequently inhibit aphid-induced calcium-mediated sieve tube occlusion in the plant, which is observed in incompatible aphid-plant interactions (Will et al., 2009). Another gene encodes the soluble calcium-activated nucleotidase 1-like isoform X2 (g12364), which has previously been annotated in whitefly salivary glands (Su et al., 2012) and is predicted to be a secretory ATP-hydrolyzing protein that could be involved in reducing the concentration of extracellular ATP and suppressing plant defenses during whitefly feeding (Roux & Steinebrunner, 2007). Altogether, these aphid DEGs and the genes discussed above (see section 2b) indicate that CaMV acquisition affects aphid saliva secretion on infected Arabidopsis. To explain this finding, we propose two non-exclusive hypotheses. In the first one, the more severe phenotype of CaMV-infected Arabidopsis, compared to TuYV-infected Arabidopsis, could induce adaptive changes of the aphid secretome to allow successful settlement on the plants. In the second hypothesis, CaMV could directly alter the saliva transcriptome. Whatever the mechanisms, these deregulations could be responsible for the changes in the feeding behavior of aphids on CaMV-infected Arabidopsis plants (Chesnais et al., 2021).

##### 3b TuYV vs CaMV in Camelina

Only 22 DEGs were found for aphids on TuYV- vs CaMV-infected Camelina, 17 expressed at higher levels in TuYV-exposed aphids and 5 expressed at higher levels in CaMV-exposed aphids (Fig. 1e). This small number of expression changes, in comparison to aphids fed on Arabidopsis, indicates strong host plant effects. They might be caused by differential host plant susceptibility to the viruses or different host-vector associations/suitability.

Among the genes expressed at higher levels in aphids on TuYV-infected vs aphids on CaMV-infected Camelina, we identified aphid genes related to development, such as the gene encoding a glycine-rich cell wall structural protein-like (g7216) implicated in chitin-based cuticle development (Table 4, see the complete list in Table S7). This again suggests that TuYV may target aphid performance by inducing morphological changes, for example, the formation of wings that could enhance transmission.

**Table 4.** Selected genes upregulated in aphids feeding on TuYV-infected vs CaMV-infected Camelina. Single lines separate genes by functional categories.

| Functional category | Potential effects on aphids | Reference(s) | Name | Gene description annotation | Top hit Taxon | Counts |  |  | log2FC | padj |
| --- | --- | --- | --- | --- | --- | --- | --- | --- | --- | --- |
|  |  |  |  |  |  | Mock | TuYV | CaMV |  |  |
| Chitin-based cuticle development | Aphid development |  | g7216 | Glycine-rich cell wall structural protein-like | *Myzus persicae* | 3672 | 4861 | 3099 | 0,65 | 1,74E-02 |
| Control of ROS and signaling | Immune system / Defense | (De Deken et al., 2014) | g9870 | Dual oxidase maturation factor 1 | *Myzus persicae* | 1699 | 2215 | 1448 | 0,61 | 2,88E-03 |
| Regulates calcium entry |  | (Hou et al., 2020) | g18794 | Calcium release-activated calcium channel protein 1-like isoform X1 | *Myzus persicae* | 1714 | 2254 | 1695 | 0,41 | 3,13E-02 |
| Hydrolase / Amino acid metabolism | Aphid metabolism |  | g24259 | Protein THEM6-like (Thioesterase-like superfamily) | *Myzus persicae* | 1060 | 1205 | 779 | 0,63 | 1,30E-02 |

Two immune-responsive aphid DEGs on Camelina were different from those observed in aphids feeding on Arabidopsis, again denoting some host specificity. One gene (g9870), expressed at higher levels in TuYV- exposed aphids than in CaMV-exposed aphids, encodes dual oxidase maturation factor 1 that is required for activation of dual oxidases and is involved in the control of reactive oxygen species (ROS) generation and signaling (De Deken et al., 2014). Its fruit fly ortholog is involved in antimicrobial defense mechanisms in the Drosophila intestine (Kim & Lee, 2014). Another gene (g18794), expressed at higher levels in TuYV-exposed compared to CaMV-exposed aphids, encodes a calcium release-activated calcium channel protein 1-like isoform X1 protein that regulates calcium entry into non-excitable cells and is required for proper immune function in Drosophila (Hou et al., 2020).

Finally, we observed that the gene coding for the protein THEM6-like (g24259) was expressed in TuYV-exposed aphids at a higher level than in CaMV-exposed aphids.

The five genes expressed at higher levels in aphids infesting CaMV-infected Camelina as compared to those infesting TuYV-infected Camelina were already discussed in previous sections of this manuscript.

Taken together, our results show that DEGs of aphids infesting TuYV-infected vs CaMV-infected Arabidopsis are quite different from those of aphids infesting another plant infected with these viruses, Camelina, even if these two plants have strong phylogenetical proximity. This reinforces the idea that responses of insect vectors had a strong host-virus specificity in our experimental system.

### Concluding remarks

We here compared the transcriptome profiles in Myzus aphids infesting two host-plant species from the family *Brassicaceae* (Arabidopsis and Camelina) infected with two viruses from different families with different transmission modes (circulative persistent TuYV and non-circulative semi-persistent CaMV). We found a strong plant species-dependent response of the aphid transcriptome to infection with either of the two viruses. This is evidenced by the higher number of aphid DEGs and stronger expression changes on virus-infected Arabidopsis compared to Camelina, regardless of the virus. Because the aphids were raised on Chinese cabbage before being transferred onto test plants for the experiments, a host switch effect might contribute to the observed transcriptome changes (Pinheiro et al., 2017; Mathers et al., 2017). However, we believe they are mostly neutralized by the experimental set-up, because the condition ‘aphid on mock-inoculated plant’ (Arabidopsis or Camelina) and not ‘aphid on Chinese cabbage’ was used as the reference for extracting mock vs virus transcriptome changes. Thus, we should have observed mostly (but probably not exclusively) changes due to viruses’ effects on aphids. It is worth noting that a plant transcriptome analysis has revealed a different picture (Chesnais, Golyaev, et al., 2022). There, TuYV altered a smaller number of plant DEGs in Arabidopsis and Camelina than did CaMV, suggesting a strong virus-specific effect on the two plant hosts. Thus, the global aphid transcriptome response to plant infection by the two viruses described here does not correlate with the global plant transcriptome response to the virus infection.

The amplitude of most expression changes was rather low (log2FC <|2|). The most obvious reason for this is technical, i.e. the use of whole aphids for RNA extraction diluted organ-specific expression changes. So, in reality, the number of DEGs and their degree of change might be higher than reported here. Only future experiments using dissected organs or micro-dissected samples will solve this issue. Nonetheless, we extracted significant information from the data. We found that stress-related aphid genes were downregulated in Myzus on both infected plants (regardless of the virus). This suggests that both CaMV and TuYV infections facilitate the establishment of Myzus on the plants, likely by downregulating expression of plant genes implicated in anti-herbivore secondary metabolism such as the jasmonic acid pathway as shown in by us in the same experimental setup (Chesnais, Golyaev, et al., 2022). Apart from common transcriptomic changes induced by both viruses, our results indicate that there are also virus-specific gene expression changes, which might be related to the transmission mode. Overall, the circulative non-propagative TuYV tended to affect developmental genes, which could increase the proportion of alate (winged) aphids in TuYV viruliferous aphids, but also contribute to their locomotion, neuronal activity and lifespan, whereas the non-circulative semi-persistent (stylet-borne) CaMV had a stronger impact on feeding-related genes and in particular those related to salivary proteins. Overall, these transcriptome alterations target pathways that seem to be particularly adapted to the transmission mode of the corresponding virus. Long-term interactions of TuYV and its aphid vectors are expected and alterations of developmental genes, potentially promoting aphid dispersion at the population level (alate morphs with higher mobility and longer lifespan), could be a suitable strategy. In support of this, we have shown increased locomotory properties of wingless TuYV-carrying aphids (Chesnais et al., 2020), but whether Myzus aphids on TuYV-infected plants also form more alate morphs remains to be shown. On the other hand, the short-term association of CaMV with the tip of the aphid stylets, together with a relatively brief retention time, should favor manipulation of rather fast processes, such as initial probing and phloem feeding, encouraging fast aphid dispersion.

Next research steps should include functional validation of the candidate genes identified in our study for their role in viral manipulation, such as aphid behavior and performance, and consequently on viral transmission. Another future research direction would be to investigate post-transcriptional changes such as post-translational protein modifications, changes in localization, metabolite composition and quantity and the like, that could likewise impact vectors but cannot be traced by transcriptomic analyses.

## Acknowledgements

We thank Claire Villeroy for aphid rearing and the experimental unit of INRAE Grand Est – Colmar (UEAV) for help with plant production and Nathalie Laboureau for technical assistance in total RNA extraction and analysis.

Preprint version 6 of this article has been peer-reviewed and recommended by Peer Community In Infections (https://doi.org/10.24072/pci.infections.100006).

## Data and supplementary information availability

Data are available online: https://doi.org/10.1101/2022.07.18.500449

The raw RNA-seq data are available under project number PRJEB54781 at the European Nucleotide Archive (https://www.ebi.ac.uk/ena/browser/view/PRJEB54781).

Supplementary information is available online: https://doi.org/10.1101/2022.07.18.500449

**Table S1.** Aligned reads for transcriptome profiling

**Table S2.** Oligonucleotides used for RT-qPCR

**Table S3.** Complete list of deregulated aphid genes in common for aphids feeding on both CaMV and TuYV-infected Arabidopsis and Camelina.

**Table S4.** Complete list of genes commonly deregulated in aphids feeding on CaMV-infected host plants (Arabidopsis and Camelina) (padj < 0.05 and log2FC > |0.5|).

**Table S5.** Complete list of genes upregulated in aphids feeding on TuYV-infected vs. CaMV-infected Arabidopsis (padj < 0.05 and log2FC > |0.5|).

**Table S6.** Complete list of genes upregulated in aphids feeding on CaMV-infected vs. TuYV-infected Arabidopsis (padj < 0.05 and log2FC > |0.5|).

**Table S7.** Complete lists of genes upregulated in aphids feeding on CaMV-infected vs. TuYV-infected Camelina and of genes upregulated in aphids feeding on TuYV-infected vs. CaMV-infected Camelina (padj < 0.05 and log2FC > |0.5|).

**Figure S1**. Quantitative reverse transcription PCR (RT-qPCR) validation of differentially expressed genes (DEGs) determined by Illumina RNA-seq profiling of the aphid transcriptome.

**Figure S2**. Kyoto Encyclopedia of Genes and Genomes (KEGG) pathways enrichment analysis of DEGs (log2FC > 1) in *Myzus persicae* in response to TuYV or CaMV infection in Arabidopsis or Camelina plants.

**Dataset S1.** RNA-seq data used to establish the heatmap.

