## Supplementary tables and figures for "Transcriptome responses of the aphid vector *Myzus persicae* are shaped by identities of the host plant and the virus": Supplementary_Tables_S1_S2_Figures_S1_S2-v5.pdf

**Table S1.** Aligned reads for transcriptome profiling

**a) Myzus on Arabidopsis**

| Sample name | Aligned reads | Uniquely mapped reads | Uniquely mapped ratio |
|---|---|---|---|
| Mp_Ara_M1 (M4) | 32690216 | 28168979 | 86,2% |
| Mp_Ara_M2 | 34814478 | 30902233 | 88,8% |
| Mp_Ara_M3 | 33434401 | 28621561 | 85,6% |
| Mp_Ara_C1 | 32344040 | 27795282 | 85,9% |
| Mp_Ara_C2 | 34954969 | 30030290 | 85,9% |
| Mp_Ara_C3 | 40965962 | 35914841 | 87,7% |
| Mp_Ara_T1 | 44442642 | 38619638 | 86,9% |
| Mp_Ara_T2 | 33535583 | 29365376 | 87,6% |
| Mp_Ara_T3 | 38100963 | 33309117 | 87,4% |

**b) Myzus on Camelina**

| Sample name | Aligned reads | Uniquely mapped reads | Uniquely mapped ratio |
|---|---|---|---|
| Mp_Cam_M1 | 32221873 | 26347102 | 81,8% |
| Mp_Cam_M3 | 30858082 | 26442426 | 85,7% |
| Mp_Cam_C1 | 34044796 | 29157873 | 85,6% |
| Mp_Cam_C2 | 34512922 | 29398003 | 85,2% |
| Mp_Cam_C3 | 38858952 | 34046290 | 87,6% |
| Mp_Cam_T1 | 36453853 | 31624484 | 86,8% |
| Mp_Cam_T2 | 41244445 | 35948989 | 87,2% |
| Mp_Cam_T3 | 32661320 | 27908031 | 85,4% |

**Table S2.** Oligonucleotides used for RT-qPCR

| Gene | Gene description annotation | Organism | Functional category | Primers |
|---|---|---|---|---|
| EF1alpha EU358933 | elongation factor-1 alpha | *Myzus persicae* | Protein synthesis | Forward primer GCCGATTGTGCTGTGCTTA<br>Reverse primer CCATCTTGTTCACACCAAC |
| g24472 | farnesol dehydrogenase-like | *Myzus persicae* | Development (Hormones metabolism) | Forward primer CAGCAGCGTGACAAAACAT<br>Reverse primer CTGCATTTCTCCGCCGACTA |
| g16389 | omega-amidase NIT2-like | *Myzus persicae* | Nitrogen compound metabolic process | Forward primer CGTCGATACCTACACGCATCA<br>Reverse primer GTCTCATCGCCTACTCGCTT |
| g22876 | calphotin-like | *Myzus persicae* | Calcium | Forward primer CGTGTCCCACAACCATACCA<br>Reverse primer TGAGGAACGTGAACTGGACG |
| g15329 | regucalcin-like isoform X1 | *Myzus persicae* | Regulation of catalytic activity, calcium binding | Forward primer CCGTGGTTCCCTATTCTCGG<br>Reverse primer GGAGATGCTCACGTTGGACA |

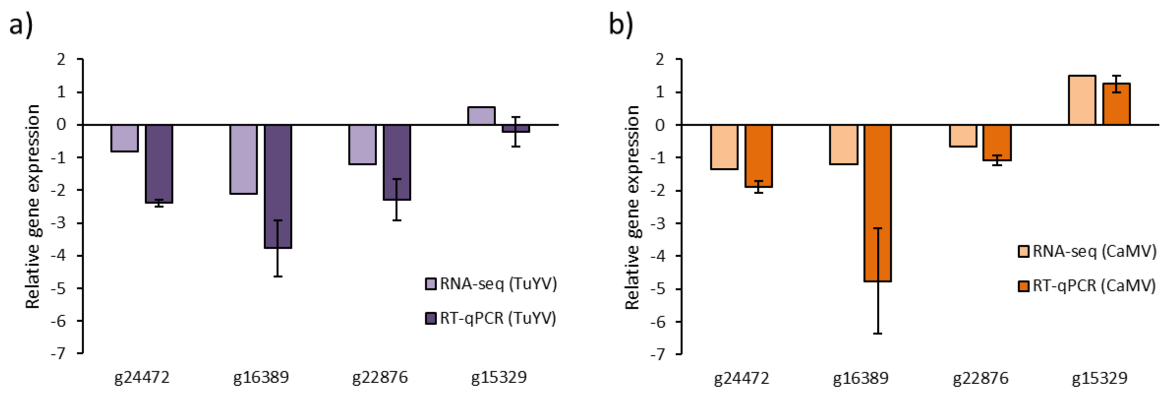


**Figure S1**. Quantitative reverse transcription PCR (RT-qPCR) validation of differentially expressed genes (DEGs) determined by Illumina RNA-seq profiling of the aphid transcriptome. a) *Myzus persicae* on CaMV-infected Arabidopsis. b) *Myzus persicae* on TuYV-infected Arabidopsis. The y-axis presents the normalized log2 fold change of expression derived from Illumina RNA-seq read counts and PCR ΔΔCp, respectively (g24472: farnesol dehydrogenase-like; g16389: omega-amidase NIT2-like; g22876: calphotin-like; g15329: regucalcin-like isoform X1).

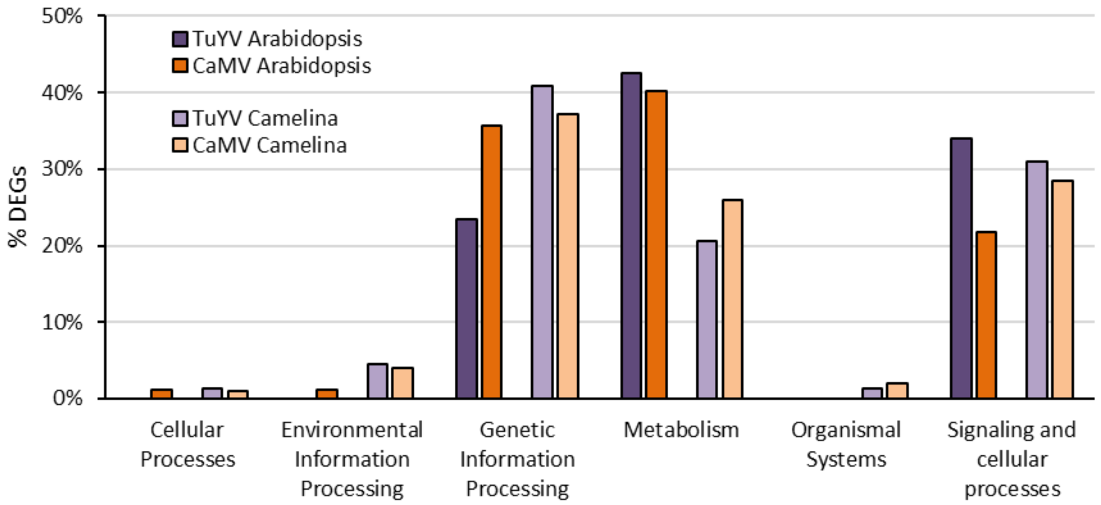


**Figure S2**. Kyoto Encyclopedia of Genes and Genomes (KEGG) pathways enrichment analysis of DEGs (log2FC > 1) in *Myzus persicae* in response to TuYV or CaMV infection in Arabidopsis or Camelina plants. Bars represent the percentage of DEGs classified into 6 main categories (Cellular Processes, Environmental Information Processing, Genetic Information Processing, Metabolism, Organismal Systems, and Signaling and Cellular Processes).
